# *In Vitro* Evolution to Increase the Titers of Difficult Bacteriophages: Rapid Appelmans Protocol

**DOI:** 10.1101/2023.03.02.530847

**Authors:** Danielle N. Kok, Joanne Turnbull, Nobuto Takeuchi, Philippos K. Tsourkas, Heather L. Hendrickson

## Abstract

Bacteriophages are becoming increasingly important in the race to find alternatives to antibiotics. Unfortunately, bacteriophages that might otherwise be useful are sometimes discarded due to low titers making them unsuitable for downstream applications. Here, we present two distinct approaches to experimentally evolve novel New Zealand *Paenibacillus larvae* bacteriophages. The first approach uses the traditional agar-overlay method, whereas the other was a Rapid Appelmans Protocol (RAP) modelled after the established Appelmans Method. Both approaches resulted in an increase in plaque-forming units (PFU/mL). The RAP approach was significantly faster and simpler, and allowed us to evolve a set of bacteriophages in as little as four days, increasing titers 100-1000-fold relative to their ancestors. The resultant titers were sufficient to extract and sequence DNA from these bacteriophages. An analysis of these phage genomes is provided. We also propose a model that describes the parameters that allow the RAP approach to select improvement of bacteriophage titer. The RAP approach is an effective method for experimentally evolving previously intractable bacteriophages in a high-throughput and expeditious manner.

## 1. Introduction

Bacteriophages (phages) are the most abundant entities on the planet, with an estimated number of 1×10^31^ virus particles, named the Hendrix product (Hendrix et al., 1999; Mushegian, 2020). The discovery, purification, and sequencing of Actinobacteriophages in undergraduate classrooms through the adoption of the Howard Hughes Medical Institute-sponsored SEA-PHAGES (Science Education Alliance Phage Hunters Advancing Genomics and Evolutionary Science) programme has brought personal empowerment through discovery to undergraduate classrooms around the globe (Hanauer et al., 2017; Russell & Hatfull, 2017). This experience has also given some the impression that bacteriophages are universally discovered and isolated with ease. The facile and frequent discovery of new Mycobacteriophages belies the lived experience of researchers who have embarked on bacteriophage hunts, only to discover that bacteriophages, even some Mycobacteriophages, are not amenable to study (LeMieux & Hatfull, 2020). One of the crucial steps in novel bacteriophage isolation is to achieve titers of over ∼5×10^9^ PFU mL^-1^, a commonly used threshold concentration of particles for electron microscopy and DNA extraction. Historically, physiological studies of bacteriophages have often been conducted at concentrations of at least 1×10^8^ PFU mL^-1^ (Delbrück, 1940; Ellis & Delbrück, 1939), although these concentrations come with their own challenges (Abedon, 2016). Achieving these high concentrations of bacteriophage particles can prove difficult for reasons that are not clear, and can subsequently derail the discovery, description, and application of newly discovered bacteriophages.

DNA sequencing is a key step in the evaluation of bacteriophages for practical use. DNA sequencing revealed aspects of the life cycle and gene content of novel bacteriophages that were not obvious by other means, including their genetic diversity and gene content, genes associated with lysogeny, host toxins, or antibiotic resistance. Genome sequences can and should influence the selection of bacteriophages for application (Philipson et al., 2018; Skurnik et al., 2007). Although genetic sequencing cannot replace *in vitro* or *in vivo* testing, it is an invaluable tool for eliminating inappropriate candidates. Complete genome sequencing of novel bacteriophages also contributes to the discovery and development of useful bacteriophage encoded enzymes and our understanding of their biology. Finally, sequencing and publishing bacteriophages furthers our understanding of the evolution of the most diverse and under sampled entities on the planet (Rohwer, 2003).

In an effort to discover bacteriophages for practical use in the apiculture industry in New Zealand, we conducted a bacteriophage hunt using the honeybee pathogen *Paenibacillus larvae. P. larvae* is a spore-forming bacterium that leads to the honey bee disease known as American Foulbrood (AFB). AFB is deleterious at both the larval and whole-hive level and can cause irreversible damage to the beehive (Elke Genersch, 2010). There have already been efforts from several laboratories around the world to try and find bacteriophages to prophylactically treat AFB, with promising results (Brady et al., 2017; Yost et al., 2016). We encountered a set of plaque-forming *P. larvae* bacteriophages that could, with effort, be brought to a titer of 3×10^7^ mL^-1^ but these proved intractable to efforts to further increase their concentrations.

A search of the literature led us to the Appelmans protocol, a 96 well-based procedure that has been used to expand the host range of bacteriophages by allowing strains to become simultaneously infected by multiple bacteriophages and allowing natural selection to screen recombinant bacteriophages for the most successful new chimaeras (Burrowes et al., 2019). Herein, we report two separate approaches we used to experimentally evolve our *P. larvae* bacteriophages, which allowed us to significantly increase their effective titer. The first approach was the experimental evolution and propagation of a bacteriophage in four parallel lineages for 25 days using a relatively low MOI of 0.05 in solid media and an agar overlay method. The second approach was a modified Appelmans protocol (Burrowes et al., 2019) that was initially performed for 30 days. We modified this approach further by allowing pure bacteriophages to adapt to single hosts rather than propagating mixed populations of bacteriophages, thereby allowing natural selection to screen mutant bacteriophages for those most able to infect the strain in liquid conditions. Subsequently, we employed this to great effect after only four days (Rapid Appelmans Protocol or RAP).

We describe both these approaches and the quality and speed of the titer improvements achieved. We report on the genomic traits of six separate bacteriophages that were evolved and subsequently sequenced using the modified Appelmans protocol. In addition, we modelled the population biology parameters operating in this brief experiment to understand the conditions under which mutation and selection act in this high-speed adaptive evolution protocol.

## 2. Materials and Methods

### 2.1 Bacterial and Bacteriophage Strains

All bacterial strains and bacteriophage isolates are listed in Table 1. Three strains of *P. larvae* were used, isolated from beehives in New Zealand with symptoms consistent with American Foulbrood disease. *P. larvae-*PaFR-2017 and *P. larvae-*PaFR-2006 were both isolated from infected honeybee larvae provided by Plant and Food Research. *P. larvae*-F1A was isolated from a symptomatic brood comb provided by AsureQuality in December 2018. The bacteriophages chosen for the 30-day protocol were all isolated from samples of soil from around healthy beehives. Phages Callan and Dash, from an apiary in the lower North Island, Phage Lilo from an apiary in the Greater Auckland region, Phage Logan from an apiary on the East Coast of North Island, Phage AJG77 from an apiary in the lower South Island and Phage ABAtENZ from an apiary in the central North Island. The bacteriophages chosen for the four-day RAP protocol were also isolated from around healthy beehives. Phage Wildcape and LunBun from an apiary in Gisborne, Phage Carlos and ApiWellbeing from an apiary in the Wellington region, Phage FutureBee from an apiary in Hamilton and Phage Rae2Bee1 from an apiary in Ashburton. Bacterial strains were grown in Brain Heart Infusion Broth (BHI) (Oxoid CM1135). Bacteriophages were grown by infecting the bacterial strain they were isolated on (see Table 1), using the agar overlay method. Bacteriophage titers were established by plating serial dilutions of the phage lysates using the agar overlay method.

**Table 1.** *P. larvae* strains and bacteriophage isolates used in this study.

| Bacterial/Phage Strain | Source | Isolation Source | Geographical Region | Notes | Accession No. |
| --- | --- | --- | --- | --- | --- |
| *P. larvae*-PaFR-2017 | Plant & Food Research | Infected larvae | Auckland |  | JARDRG000000000 |
| *P. larvae*-PaFR-2006 | Plant & Food Research | Infected larvae | Hamilton |  | JARDAI000000000 |
| *P. larvae*-F1A | AsureQuality | Symptomatic Brood Frame | North Canterbury |  | JARDRL000000000 |
| Phage Callan | This work | Soil | Wellington | Isolated on PI-PaFR-2006 | OP503989 |
| Phage Dash | This work | Wax | Wellington | Isolated on PI-PaFR-2006 | OP503990 |
| Phage Lilo | This work | Soil | Waikato | Isolated on PI-F1A | OP503991 |
| Phage Logan | This work | Soil | Gisborne | Isolated on PI-PaFR-2017 | OP503980 |
| Phage AJG77 | This work | Soil | Otago | Isolated on PI-PaFR-2017 | OP503969 |
| Phage ABAtENZ | This work | Soil | Waikato | Isolated on PI-PaFR-2017 | OP503968 |
| Phage Wildcape | This work | Soil | Gisborne | Isolated on PI-F1A | OP503988 |
| Phage Carlos | This work | Soil | Wellington | Isolated on PI-F1A | OP503973 |
| Phage ApiWellbeing | This work | Soil | Wellington | Isolated on PI-F1A | OP503970 |
| Phage LunBun | This work | Soil | Gisborne | Isolated on PI-F1A | OP494865 |
| Phage FutureBee | This work | Soil | Waikato | Isolated on PI-TP | OP503975 |
| Phage Rae2Bee1 | This work | Soil | Ashburton | Isolated on PI-TP | OP503983 |

### 2.2 Experimental evolution to increase the infectivity of P. larvae bacteriophage Lilo

Multiplicity of infection (MOI) was established by plating serial dilutions of both colony-forming units (CFU) and plaque-forming units (PFU). Specifically, a single colony isolate of the *P. larvae* strain was inoculated in 5 mL of BHI broth, incubated at 37°C and shaken at 100 rpm for 2 days. This *P. larvae* liquid culture was serially diluted to a total dilution factor of 10^−8^ and 5 µL of each dilution was applied to 1.5% agar BHI plates. Plates were incubated without shaking at 37°C for two days. Colonies were counted to determine the number of CFU per mL of culture. PFU determination was similar, the *P. larvae* bacteriophage Lilo lysate was serially diluted to a factor of 10^−8^ and 5 µL of each dilution was applied as a spot test to a *P. larvae* bacterial lawn plated in 0.5% BHI top agar. The plate was incubated without shaking at 37°C for two days. The number of plaques observed for each dilution was counted to estimate the titer of the lysate.

Four biological replicates of bacteriophage Lilo were serially passaged at a low MOI. At each serial passage, 500 µL of bacterial culture was inoculated with lysate for an estimated MOI of 0.05 (2×10^6^ PFU were plated with ∼4×10^7^ CFU). These were incubated without shaking at room temperature for 30 minutes to facilitate adhesion before they were plated in 3 mL of 0.5% top agar BHI with 1 mM CaCl_2_, 1 mM MgCl_2_ and 1mM thiamine hydrochloride, onto 1.5% agar BHI plates. Plates were incubated without shaking at 37°C for two days. The confluent or cleared plates were flooded with 8 mL BHI broth and incubated without shaking at room temperature for 2-4 hours. The resulting lysate was collected and passed through a 0.45 µm syringe filter, in preparation for use in the next passage. Lysates from each passage were titered by spot plate assay. Plaque counts were used to determine lysate titers from each passage and used to adjust the volumes plated in subsequent passages to maintain the desired MOI of 0.05. Plaque diameters were measured using ImageJ software (Abràmoff & Magalhães, n.d.).

### 2.3 Modified Appelmans Protocol in 96-well plates

We implemented the modified Appelmans protocol as in Burrowes *et al*. with changes to the media and by restricting the horizontal wells in the 96-well plate to a single strain of host and a single strain of bacteriophage (Burrowes et al., 2019). Using a 96-well plate, we seeded 100 µL of serially diluted bacteriophage lysate in 100 µL double-strength BHI Broth containing 4 µL culture of a single strain (Fig. 2a). Plates were incubated at 37°C on a shaking platform at 100 rpm. Wells showing complete lysis, plus the first turbid well, were pooled. If no lysis was observed, wells containing undiluted bacteriophage were harvested. The pooled lysates were sterilised by vortex mixing with 1:100 CHCl_3,_ left for 10 minutes and subsequently centrifuged at 15,000g for 15 minutes. Lysates were removed, leaving the CHCl_3_, and transferred to fresh tubes for storage. These pooled lysates were used to initiate the next round of directed evolution in a similar manner. Pooled lysates were tested for titer after every second round using the agar overlay spot titer method.

### 2.4 Genomic DNA Extraction

Evolved bacteriophages were triple purified by selecting a single plaque from a plate and using this to infect in a standard agar overlay. Bacteriophages from the final round of purification were used to inoculate a 60 mL culture of the bacterial strain they were isolated on at a multiplicity of infection (MOI) of 0.1. Cultures were incubated at 37°C with shaking at 100 rpm overnight. Bacteriophage lysates were purified as described previously (Bonilla et al., 2016). Briefly, an aliquot of bacteriophage lysate was transferred into a falcon tube and centrifuged at 4,000 × g for 20 min, the supernatant was collected and transferred into a new falcon tube. The lysate was filter-sterilised through a 0.22 µm filter, and chloroform was added to bring the total volume of the lysate to 1.10 (0.1 of volume in chloroform). The lysate was then vortexed and incubated at room temperature for 10 min. The lysate was then centrifuged at 4,000 × g for 5 min and transferred into Nalgene Oak Ridge High-Speed PPCO Centrifuge Tubes, leaving the chloroform behind. The sterilised bacteriophage lysate was then concentrated by centrifugation at 18,000 rpm for 45 minutes. Subsequently, 50 mL of this pelleted lysate was re-suspended into a 5 mL volume of BHI. The bacteriophage DNA was extracted using a modified zinc chloride precipitation method (Santos, 1991). Modifications included the addition of 1 µL Proteinase K which was incubated at 37°C for 10 min after the TES step. The tubes were left on ice overnight after the isopropanol was added. On day 2 of the protocol 1 µL of pure glycogen was added to each tube before centrifugation to aid in pelleting and visualisation of the DNA. DNA pellets were resuspended in 50 µL nuclease-free water.

### 2.5 Library Preparation and Sequencing

A total amount of 1µg DNA per sample was used as input material for the DNA sample preparations. Sequencing libraries were generated using NEBNext® Ultra™ DNA Library Prep Kit for Illumina (NEB, USA) following the manufacturer’s recommendations and index codes were added to attribute sequences to each sample. Briefly, the DNA sample was fragmented by sonication to a size of 300 bp. DNA fragments were end-polished, A-tailed, and ligated with the full-length adaptor for Illumina sequencing with further PCR amplification. At last, PCR products were purified (AMPure XP system) and libraries were analysed for size distribution by Agilent2100 Bioanalyzer and quantified using real-time PCR. The clustering of the index-coded samples was performed on a cBot Cluster Generation System according to the manufacturer’s instructions. After cluster generation, the library preparations were sequenced on an Illumina NovaSeq 6000 platform and paired-end reads were generated.

### 2.6 Genome Assembly

Bacteriophage genome assembly was carried out using Geneious 9.05 (Auckland, New Zealand) (https://www.geneious.com), using the built-in Geneious assembler, with either Medium Sensitivity/Fast or Medium-Low Sensitivity/Fast, and circularise contigs with matching ends selected.

The genome ends and DNA packaging strategy were identified by sequence similarity to already published *P. larvae* bacteriophages (Stamereilers et al., 2018). Bacteriophages were searched for the two known 3′ overhang sequences “CGACGGACC” or “CGACTGCCC” near the terminase genes (Stamereilers et al., 2018). Phages AJG77, ABAtENZ and Logan were found to have the “CGACGGACC” sequence, while Dash, Callan and Lilo contained the “CGACTGCCC” sequence. Genomes were rearranged so these 3′ overhang sequences were at the end of the genome. This resulted in the small terminase gene starting at either 50 or 51 base pairs downstream of base 1. Genes were identified by running the rearranged files through Phage Commander (Lazeroff et al., 2021) with all gene identification programs selected. The GenBank-formatted output files were entered in DNA Master (cobamide2.bio.pitt.edu) to manually check for false positives, missing genes and identify start codons as described in detail in (Salisbury & Tsourkas, 2019). Putative protein functions were assigned as described in previous work (Stamereilers et al., 2018).

### 2.7 Host Range Assays

The ability of bacteriophages to infect an isolate was assessed by 3 µl spots of a bacteriophage lysate onto a double-layer agar containing 500 µl of bacterial lawn. Each *P. larvae* bacterial isolate was tested separately.

### 2.8 Simulation to Model Mutation Frequency in Phages

We constructed a mathematical model to estimate the number of mutations that could develop in this RAP experiment. The numbers of available cells, phages and time have been input from the experimental protocol. The model to simulate the number of phage mutants at the end of 4 days of experimental evolution assumes the following kinetic equations:

The first equation represents the infection of a bacterium by a phage:

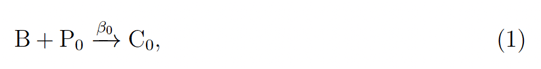

where B is a bacterium, P_0_ is a phage with genotype 0, *β*_0_ is the infectivity of P_0_, and C_0_ is a bacterium infected by P_0_.

The second equation represents the lysis of an infected bacterium:

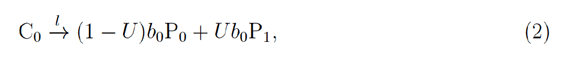

where *l* is the rate of lysis, U is the mutation rate per replication per genome, *b*_0_ is the burst size of P_0_, and P_1_ is a phage with genotype 1. The value of *b*_0_ is assumed to be constant.

There are two more equations, which describe the infection and lysis by P_1_:

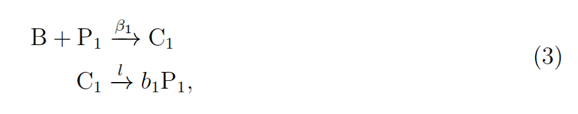

where *β*_1_ is the infectivity of P_1_, and *b*_1_ is the burst size of P_1_.

We assume that *b*_1_ = *b*_0_(1+*s*_*b*_) and *β*_1_ = *β*_0_(1+*s*_*β*_), where *s*_*b*_ and *s*_*β*_ are the selection coefficients. For example, if *s*_*b*_ = 1, the burst size of P_1_ is twice that of P_0_.

For simplicity, we do not assume the spontaneous decay of phages or that there are more than two genotypes of phage. The above kinetic equations are simulated with the Gillespie algorithm. The model assumes 10 tubes, each containing 40,000 B at the beginning as in the experimental evolution described above.

At the start of each simulation, the initial numbers of P_0_ in different tubes are set to numbers drawn from Poisson distributions with the following means: 1,000 (zeroth tube), 100 (first tube), 10 (second tube), etc. Thus, about three tubes will have any phage particles at the beginning of Day 0. The initial number of P_1_ is set to zero in all tubes. The above initial condition is equivalent to assuming that the concentration of phage particles obtained from the solid medium before the experimental evolution is ten phage particles per µL.

All tubes are incubated until all bacteria (B) are lysed in the tubes that have at least one phage particle P_0_ or P_1_.

The tubes that have at least one phage particle are pooled, plus one tube that has no phage particles. All tubes are assumed to contain 200 µL of media. 100 µL of the pooled media is added to the zeroth tube, 10 µL to the first tube, 1 µL to the second tube, etc. The number of phage particles in each tube is again determined by drawing a number from a Poisson distribution with the corresponding mean number of phage particles.

The tubes are incubated again until all bacteria are lysed in the tubes that have at least one phage particle. In a typical simulation, about five tubes contain phages from Day 1. The above repeats up to three transfers, i.e., four days.

One hundred simulations were run for each condition tested. Any arbitrary large value could be used, we also spot-checked running the model with one thousand simulations with similar results. Parameters were varied within experimentally reasonable values as described in the results.

## 3. Results

Using strains of *P. larvae* as hosts, 26 novel *P. larvae* bacteriophages were isolated using standard bacteriophage discovery methods. Briefly, direct isolation and enrichment methods were employed and bacteriophages able to form plaques were discovered and isolated from hive materials, bee debris and soil from around or within healthy, non-infected hives. The complete process of the discovery of these *P. larvae* bacteriophages will be described elsewhere (Kok, Zhou, and Hendrickson *in preparation*).

The plaques of the six bacteriophages used in the 30-day modified Appelmans Protocol were pinprick size plaques in 0.5% top agar. We employed a suite of standard methods for increasing the titers of these bacteriophage lysates, but none were successful (webbed plates, liquid infections, large volume infections and enhanced centrifugation). The titers of each lysate generally remained between 5 × 10^5^ mL^-1^ or up to 3 × 10^7^ mL^-1^, 1,000-10,000 times too low for DNA extraction or electron microscopy.

Unable to proceed further in our characterisation, we chose to experimentally evolve these bacteriophages in the hopes that by selecting for increased efficiency in host infection we would be able to increase the titers sufficiently to carry on with our studies of these novel bacteriophages. We first employed a method that used an agar overlay, reasoning that we had observed plaques and had not observed success in propagating these bacteriophages in liquid.

### 3.1 Agar Overlay Method of Evolution Improves Titer

We developed an experimental protocol in which bacteriophages were serially propagated in top agar overlays with an abundance of host bacteria. We chose phage Lilo and performed infections using a relatively low MOI of 0.05 (1 bacteriophage for 20 bacterial cells) reasoning that bacteriophages capable of rapid infection and higher burst sizes would be favoured. Four low MOI Lilo lysates were continually harvested and propagated on the ancestral bacterial strains for 25 days (Fig. 1a). Bacteriophage titers were calculated after each overnight passage for all Lilo replicate lines (Lilo E, F, G and H). From the initial titer of 3 × 10^7^ PFU mL^-1^ for the starting lysate, after 15 passages an increase in titer of 4.1 to 14.5-fold was observed in three out of four lines (Fig. 1b, n = 1). Those three lines maintained an average 5 to 7-fold increase from the starting lysate’s titer. Lilo E maintained an average increase of 3-fold throughout the experiment.

**Figure 1.**
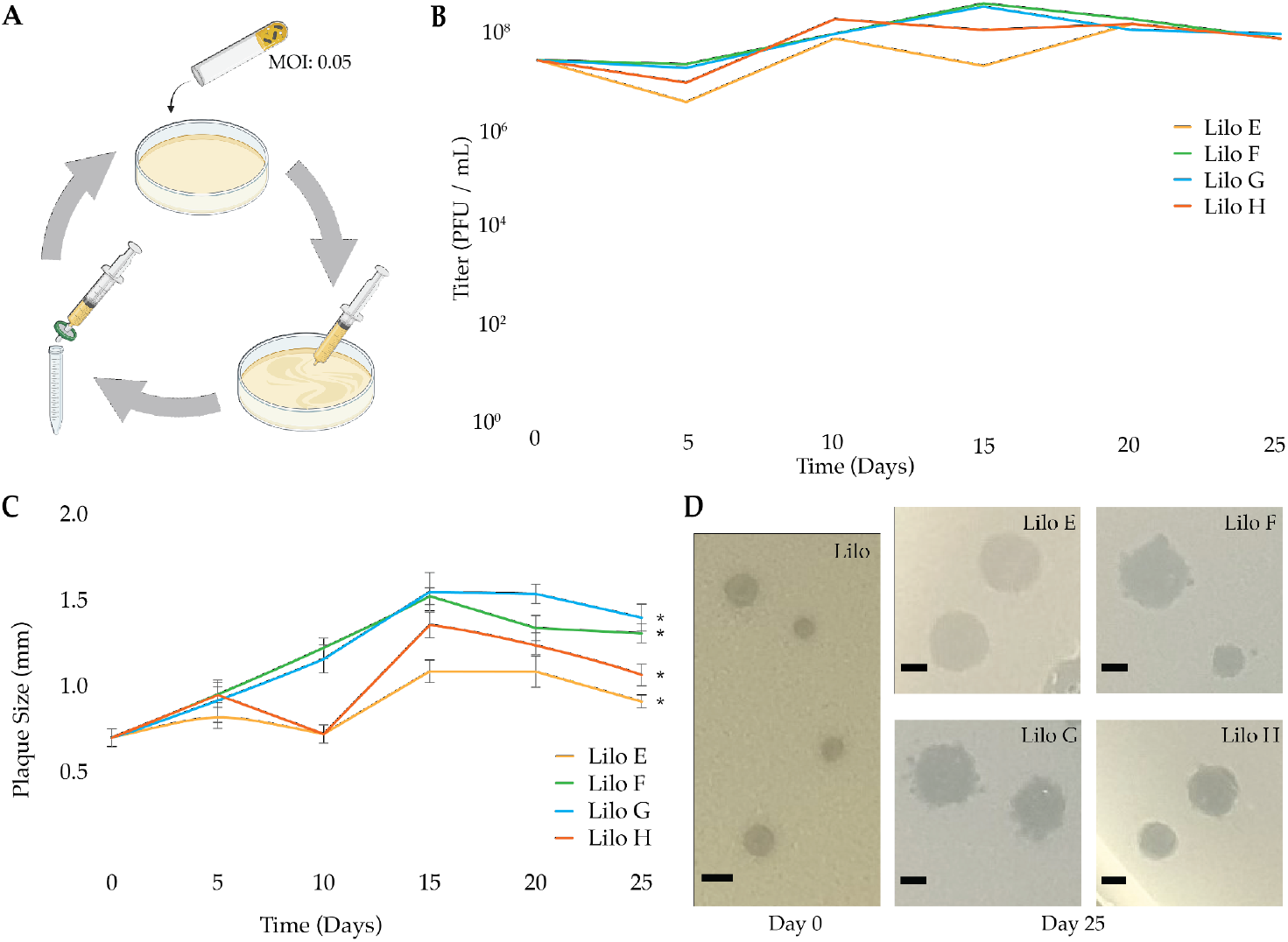
Experimental evolution of bacteriophage Lilo on solid media. (**a**) Schematic of experimental method (created with BioRender.com). (**b**) Increase in bacteriophage titer (PFU mL^-1^) over the 25-day experimental evolution for four bacteriophage Lilo lineages E, F, G and H. (**c**) Increase in plaque diameter from the initial bacteriophage lysate until day 25 of the evolution experiment, (\**P* <0.05; paired *t*-test). Error bars = standard error. (**d**) Representative plaques for the ancestral Lilo and Lilo lineages E, F, G and H after 25 days. Scale bar = 1 mm.

**Figure 2.**
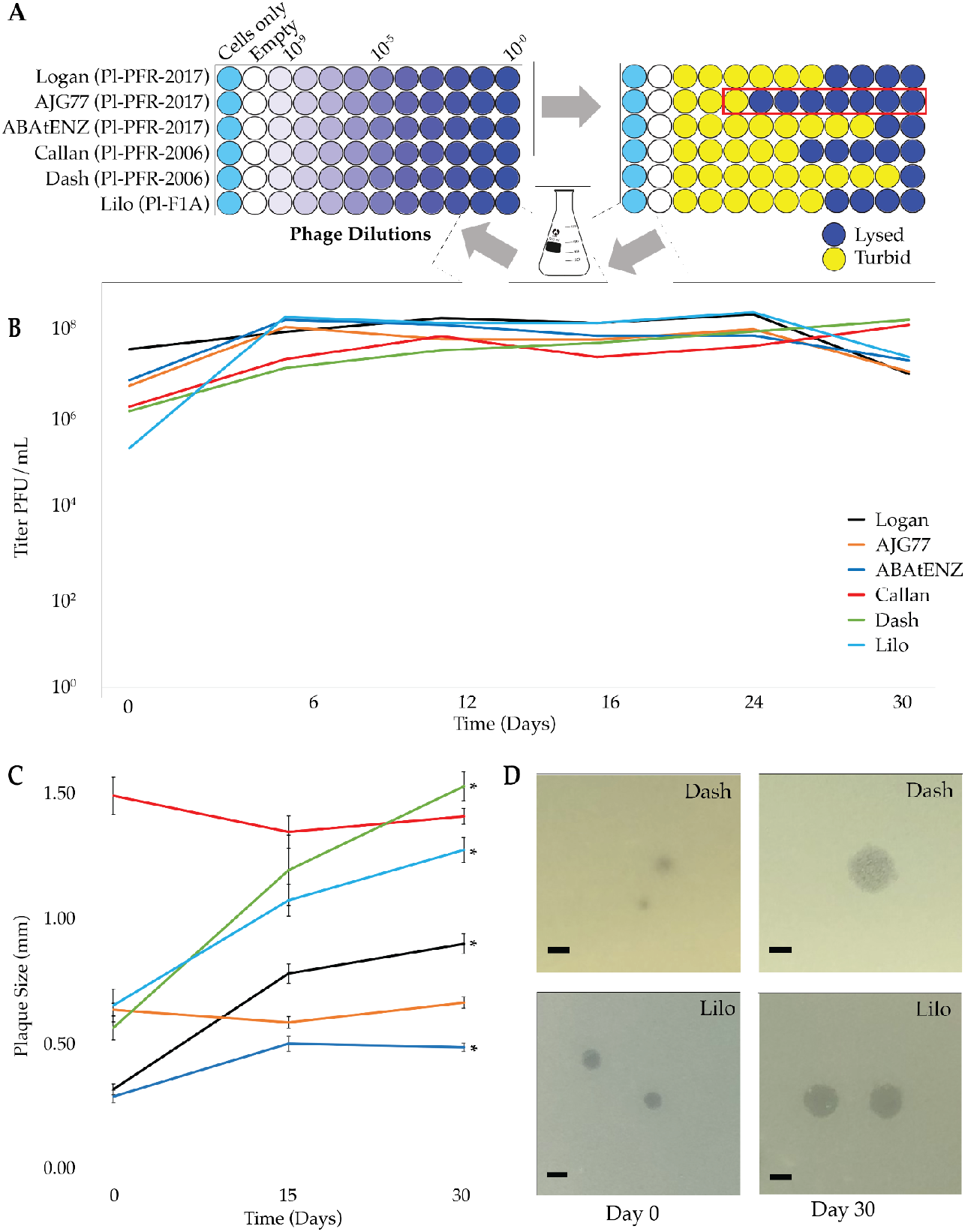
Experimental evolution using a modified Appelmans Technique. (**a**) Schematic of 96-well format modified Appelmans protocol. (**b)** Increase in bacteriophage titer (PFU/mL) over the 30-day experimental evolution for six bacteriophages Logan, AJG77, ABAtENZ, Callan, Dash and Lilo. (**c**) Increase in plaque size from the initial bacteriophage lysate until day 30 of the evolution experiment, (\**P* < 0.05; paired *t*-test). Error bars = standard error (**d**) Change in plaque size over 30 days for phage Dash and Lilo. Scale bar = 1 mm.

### 3.2 Plaque Improvements from Agar Overlay Method

Plaque size changes can indicate a mutation has taken place in a bacteriophage of interest (Gallet et al., 2011). The ancestral Lilo bacteriophages created plaque diameters that measured at 0.70 mm (+/-0.05) (mean, n = 10). In order to assess whether the experimental evolution was having an effect on the bacteriophages, diameters of the evolved Lilo plaques were measured after each overnight passage. After 15 passages a 56% -121% increase in the mean diameter of plaques was observed in all four evolved Lilo lines, with lines E-H measuring an average of 1.09 (+/-0.06) mm, 1.52 (+/-0.04) mm,

1.55 (+/-0.11) mm and 1.36 (+/-0.07) mm respectively (Fig. 1c). The increase in plaque diameter was maintained throughout the remainder of the 25-day experiment, however, some of these apparent gains were not retained in the course of the selection. Lines F and G saw the greatest increase in plaque size (Fig. 1d), measuring 1.31 (+/-0.05) mm and 1.40 (+/-0.07) mm respectively at the end of the 25 days, while lines E and H measured 0.91 (+/-0.04) mm and 1.06 (+/-0.06) mm respectively (Fig. 1c). Ultimately, the maximum increase in plaque size at the end of the experiment was Lilo G (1.40 (+/-0.07) which doubled from the initial Lilo plaque size of 0.70 (+/-0.05) mm.

### 3.3 Modified Appelmans Protocol to Improve Bacteriophage Titers

Whilst the top agar overlay evolution experiment showed signs that mutations had likely taken place, the increases in titer achieved (4-14-fold) were not sufficient to justify the effort of propagating bacteriophages in top agar overlays for this period of time. We, therefore, turned our attention to the literature and found the Appelmans protocol described by Burrowes and colleagues (Burrowes et al., 2019). The Appelmans protocol is a small volume, high throughput directed evolution experiment administered to foster recombination between similar bacteriophages in liquid lysates over a period of 30 days. Historically, this method has been used to increase host range by passaging serial dilutions of mixed bacteriophages on several hosts in a 96-well plate format (Fig. 2a).

We adopted the basic protocol in order to use this passaging procedure to apply selection pressure on populations of a *single* bacteriophage on a *single* host to improve infection properties in liquid media over a similar time period (Fig. 2a). This modified Appelmans protocol was performed on six novel *P. larvae* bacteriophages. Bacteriophage efficacy was checked by measuring the PFU after every second transfer (Fig. 2b). Initial bacteriophage titers of our 6 bacteriophages ranged from 2 × 10^5^ mL^-1^ -3 × 10^7^ mL^-1^. After 30 days of the modified Appelmans Protocol the titers of these bacteriophages had increased to between 1 × 10^7^ mL^-1^ -2 × 10^8^ mL^-1^, a 100-fold increase on average (Fig. 2b).

### 3.4 Plaque Improvements from Modified Appelmans Protocol

A significant increase in plaque size was observed in four out of the six bacteriophages. Plaque sizes were measured on Day 0, Day 15 and Day 30 (Fig. 2c). AJG77 and Callan did not exhibit larger plaque sizes and measured approximately 0.65 (+/-0.02) mm and 1.40 (+/-0.05) mm respectively for the duration of the experiment. ABAtENZ, Logan, Lilo and Dash increased in their respective plaque sizes from 0.31 (+/-0.02) mm, 0.33 (+/-0.02) mm, 0.66 (+/-0.06) mm and 0.57 (+/-0.04) mm to 0.50 (+/-0.01) mm, 0.90 (+/-0.03) mm, 1.27 (+/-0.04) mm and 1.52 (+/-0.09) mm respectively (*P* <0.05; paired *t*-test). ABAtENZ and Lilo (Fig. 2d) achieved 63% and 92% increases in plaque size respectively over 30 days while Dash (Fig. 2d) and Logan saw the greatest increase in plaque size with an increase of approximately 165% and 170% respectively.

### 3.5 Similar Results Seen After Four Days

The original Appelmans protocol used a 30-day time period. The Modified Appelmans Protocol experiment described above appeared to yield significant increases in bacteriophage titers after the first few rounds of plating (Fig. 2b). This was particularly evident in Lilo. We, therefore, selected a new set of six novel *P. larvae* bacteriophages and subjected them to a 4-day Rapid Appelmans Protocol (RAP) treatment (Fig. 3). The initial titers of the six bacteriophages selected for RAP ranged from 1 × 10^4^ mL^-1^ -1 × 10^5^ mL^-1^, with Rae2Bee1 having the lowest titer and Wildcape the highest titer. After four days of RAP evolution, the titers increased to 2 × 10^7^ mL^-1^ -4 × 10^8^ mL^-1^ a 1000-fold increase on average (Fig. 3a). Rae2Bee1 showed the smallest increase in titer, with a 220-fold increase, ApiWellbeing, Carlos, and Wildcape all experienced a 1000-fold increase in titer, and LunBun and FutureBee saw the largest increases in titer with increases of 2,700 fold respectively. An increase in plaque size was also seen in all six bacteriophages ranging from an increase of 53% to 514%, FutureBee and Carlos respectively (Fig. 3b). Interestingly, the average increase in plaque size did not correlate with increases in titer as FutureBee experienced the largest increase in titer but the smallest increase in plaque size. Ultimately, we applied either the RAP or the longer protocol to all 26 of our *P. larvae* phages. We observed there is a correlation between the starting titer and the titer improvement which suggests that there is a limit to the titer enhancement that can be achieved in the RAP protocol (Fig. 3C).

**Figure 3.**
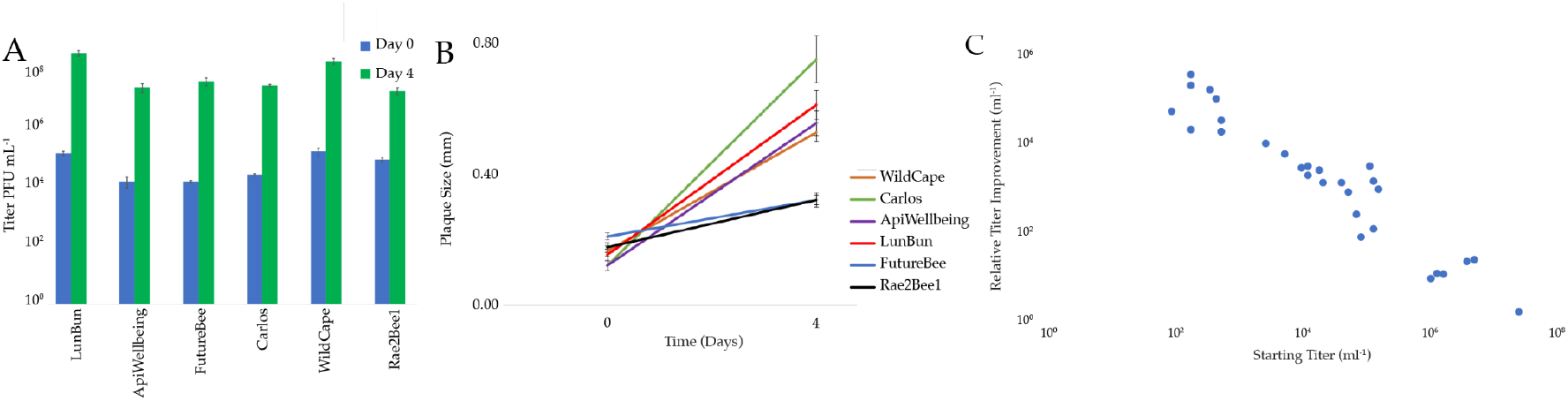
RAP Experimental evolution increases lysate titer in as little as four days. (**a**) Increase in bacteriophage titer (PFU/mL) in four days for six bacteriophages. Error bars = standard error (N=3). (**b**) Colony measurements for the RAP evolved bacteriophages in A. **(c)** The RAP protocol appears to limit the potential for phages to improve based on their starting titer.

### 3.6 Appelmans Leads to Bacteriophage Genomes

The resulting lysate concentrations after both modified Appelmans methods were sufficient to extract high-quality DNA for Illumina sequencing. Sequencing results are shown for phages that underwent the longer 30-day protocol (Table 2 and Fig. 4). The genomes of these phages ranged between 40kbp and 44kbp in length and have 70-82 genes each. Pairwise comparisons of these genomes were carried out using Phamerator, a bioinformatics tool that uses the “Align Two Sequences” program contained within BLAST (Cresawn et al., 2011). The maps show how related or divergent the phages are, showing that Lilo, Callan, and Dash are highly related, whereas ABAtENZ, AJG77, and Logan are highly related to each other but diverge from the other three phages apart from two regions of the structural genes (Fig. 4). The details of these genomes and their functional genes will be discussed elsewhere (Kok, Tsourkas, Gosselin, and Hendrickson *in preparation*).

**Table 2.** The genomic characteristics of the first six bacteriophage genomes that we sequenced using the Modified Appelmans Protocol to increase titers.

|  | Genome Length (bp) | DNA Packaging Strategy | GC Content (%) | No. of Genes | Genes per 1000bp | Percent Coding (%) |
| --- | --- | --- | --- | --- | --- | --- |
| **Logan** | 44,419 | 3' cos | 43.0 | 82 | 1.85 | 94.01 |
| **AJG77** | 44,417 | 3' cos | 43.0 | 82 | 1.85 | 93.46 |
| **ABAtENZ** | 44,419 | 3' cos | 43.0 | 82 | 1.85 | 94.01 |
| **Lilo** | 40,941 | 3' cos | 40.3 | 70 | 1.71 | 91.78 |
| **Callan** | 44,768 | 3' cos | 39.6 | 77 | 1.72 | 91.56 |
| **Dash** | 44,599 | 3' cos | 39.4 | 79 | 1.77 | 93.40 |

**Figure 4.**
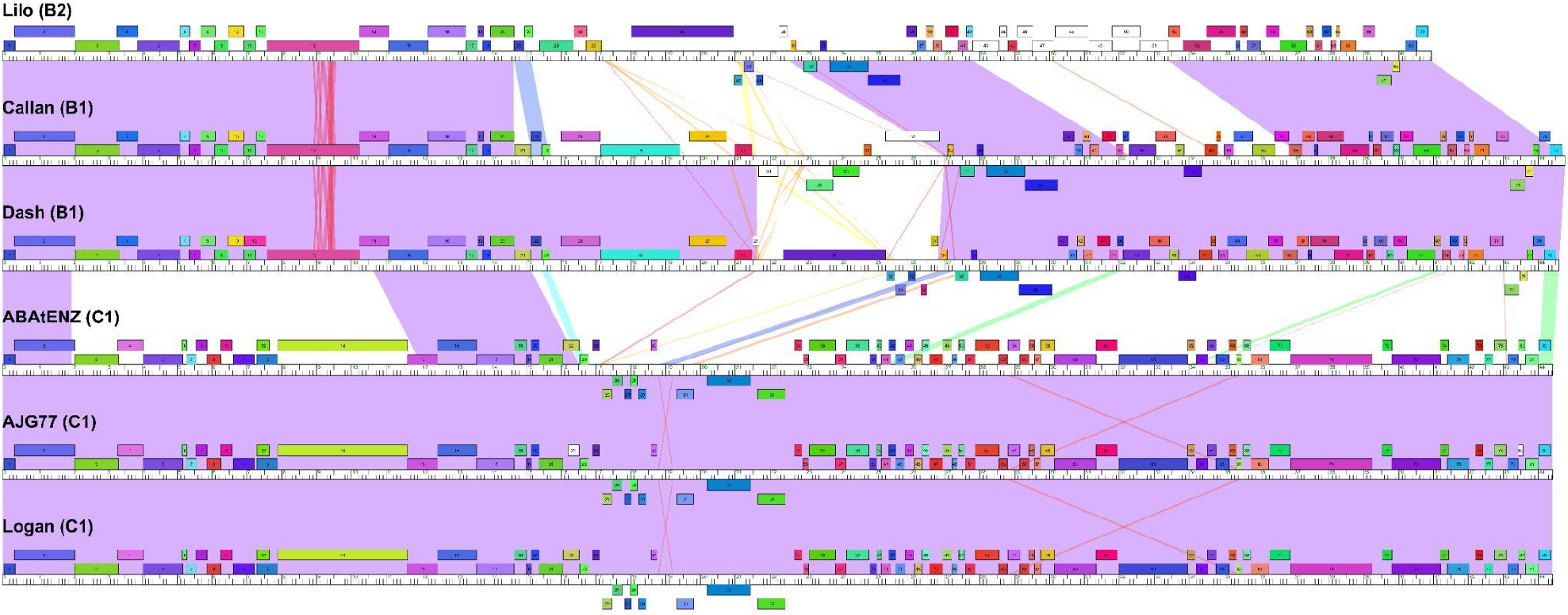
Pairwise genome maps of bacteriophages evolved for 30 days. Maps generated by Phamerator. Shading indicates high sequence similarity between sequences as determined by BLASTN, with purple being the highest. (E-value = 0). Genes are represented by boxes, boxes with the same colour indicate genes which belong to the same families (“phams”).

### 3.7 Changes to Host Range of New Zealand P. larvae phages after Appelmans protocol

The host range of all 26 *P. larvae* phages was assessed against 30 New Zealand *P. larvae* bacterial strains before they underwent the Appelmans protocol. The host range was then checked again after the phages had been evolved. An expansion in the host range was seen in 32 instances (red asterisks in Fig. 5) in 11 phages, and there were no instances of a reduction in the host range. Eight instances of expanded host range occurred in bacterial strain W19_08100, while five instances occurred in W19_08099, both of which had a narrower host range than the majority of bacterial strains before the evolution of the phages. ABAtENZ gained the ability to infect an additional five bacterial strains, which were only able to be infected by Callan, Dash, and Lilo before the Appelmans protocol. As we were unable to sequence these phages before evolution, we could not compare their genomes before and after evolution to ascertain what may have caused the expansion of the host range.

**Figure 5.**
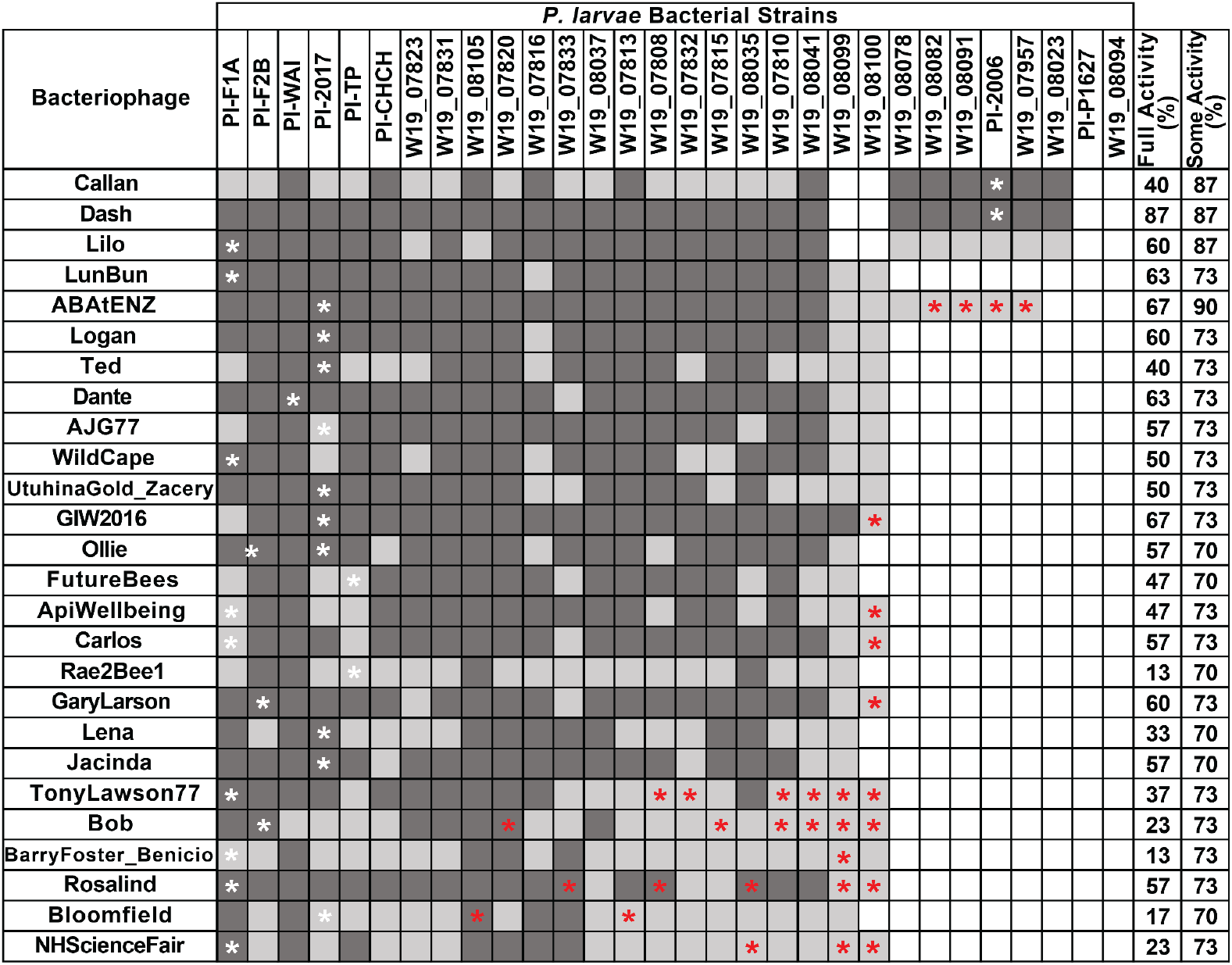
Host range of 26 *P. larvae* bacteriophages on 30 *P. larvae* bacterial isolates from New Zealand. Dark grey boxes indicate complete cell lysis, light grey boxes indicate some cell lysis and white boxes indicate no cell lysis has occurred. Red asterisks show where host range expansion has occurred. White asterisks show the original *P. larvae* strain the phage was isolated on.

We investigated the possibility that mutations might have occurred that allowed escape from CRISPR recognition. Previously sequenced *P. larvae* strains are known to have CRISPR-Cas systems, and *P. larvae* phages may have evolved point mutations to escape their hosts’ CRISPR systems (Stamereilers et al., 2021). We subjected preliminary sequences (in contigs) of our eight *P. larvae* strains to CRISPRFinder (Grissa et al., 2007) in order to identify 38 unique CRISPR spacers.

We searched the phage genomes for these *P. larvae* CRISPR spacers allowing up to 80% nucleotide divergence (approximately 7bp changes allowed). Callan, Dash, and Lilo all contain spacers that range from 81% to 100% nucleotide identity to at least one *P. larvae* CRISPR spacer found in our sequenced isolates. However, these three phages did not experience any expansion in their host range after the Appelmans protocol. Based on this analysis, we cannot determine what the mechanism of host range expansion was but it does not appear to be caused by the evasion of CRISPR spacers.

### 3.8 Modelling of Rapid Appelmans Protocol

To model the possibility of a beneficial mutation becoming dominant after as little as four days of evolution in the RAP we assembled a simulation. The initial parameters for the simulation were as follows:

- Rate of lysis: *l* = 1/30 [min^−1^], meaning that lysis takes, on average, 30 minutes
- Basal infectivity: *β*_0_ = 10^−6^ [virion^−1^ min^−1^]
- Basal burst size: *b* = 10

Under these parameters it would take approximately 10 hours for all 40,000 bacteria to be lysed. The parameters U (mutation opportunity), *s*_*b*_ (burst size coefficient), and *s*_*β*_ (infectivity coefficient) were varied as follows:

Mutation opportunity (U) was set between 0.0 × 10^0^ to 1.0 × 10^−5^. This is equivalent to multiplying the mutation rate, 5 × 10^−7^ per base pair (or Drakes rule (Drake, 1991)), by the number of possible sites that could be mutated (out of ∼40,000 bp) that would result in an increase in the burst size of that phage. We set this second part of the U parameter to between 0 and 20 nucleotide sites (eg. 20 × 5 × 10^−7^ = 1.0 × 10^−5^). The selection coefficient of burst size (*s*_*b*_) was set between 1 and 60 (meaning that a mutation increases the burst size by 1 to 60-fold). The selection coefficient of infectivity (*s*_*β*_) was set between 1 to 10-fold increase in infectivity. This means that depending on the parameters set for any given simulation the largest potential increase in total fitness due to a single mutation would result if an increase in burst size of 60-fold and an increase in infectivity of 10 fold were chosen, e.g. 60 × 10 = 600 fold increase in phage fitness.

One hundred simulations were run for each chosen set of parameters and a frequency histogram of the fraction of the mutant phage after 4 days was obtained. Histograms showed a bimodal distribution, showing peaks at 0 and 1 (Figure 6). This bimodality comes from the fact that the limiting step for the mutant phage to emerge in a population is the occurrence of a fitness-improving mutation. If the mutant occurs, it can spread within a small amount of time. Figure 7 shows the percentage of experiments that resulted in populations in which 70% or more of the phages in the simulation were mutants with a fitness advantage after four days of RAP. Setting the mutation opportunity (U) to 0 resulted in zero mutant phages regardless of parameters affecting increased burst size or infectivity.

**Figure 6.**
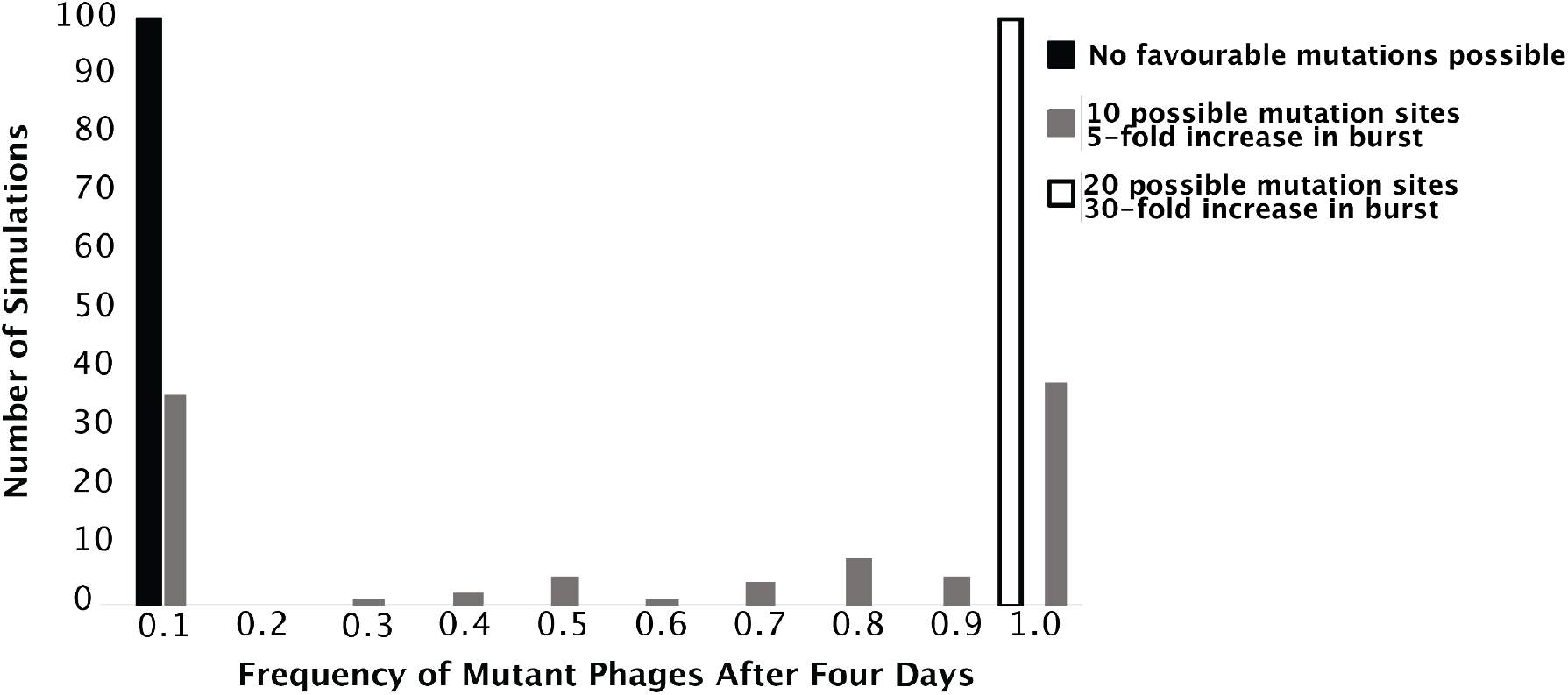
Histogram of three representative simulations when infectivity is set to one. Black shows when parameters are set so the simulation results in zero mutant phages (U = 0, *s*_*b*_ = 1, *s*_*β*_ = 1). Red shows the transition point or at least 50% of the experiments results in 70% or more of the phages having the fitness-increasing mutation (U = 5.0 × 10^−6^, *s*_*b*_ = 5, *s*_*β*_ = 1). Blue represents the upper limit of the simulation or when all experiments have at least 70% fitness-increasing mutant phage after four days (U = 1.0 × 10^−5^, *s*_*b*_ = 30, *s*_*β*_ = 1).

**Figure 7.**
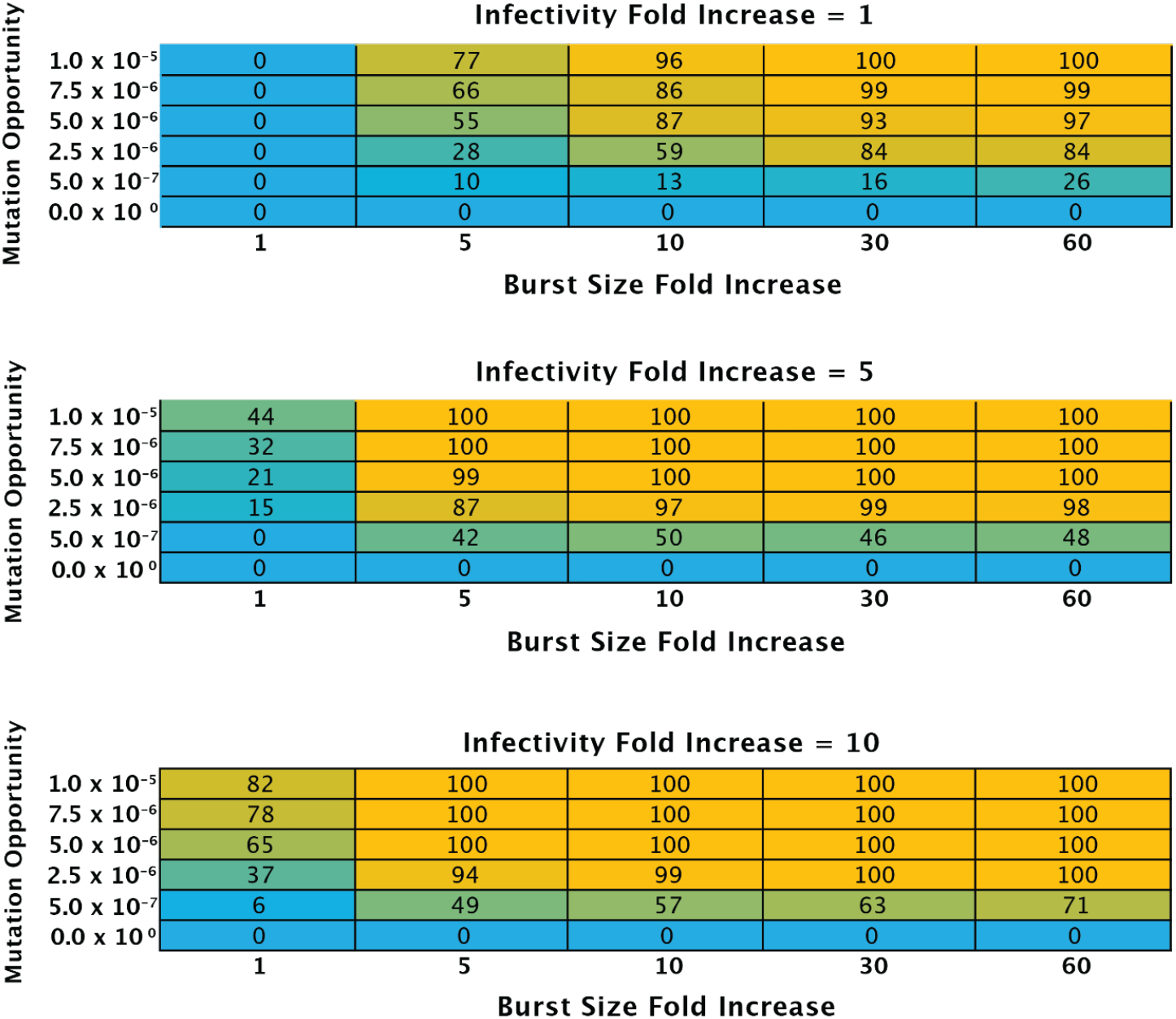
Heat maps of simulations of RAP. One hundred simulations were run for each change in parameters. Mutation opportunity, increase in burst size, and increase in infectivity coefficient were varied as follows: Blue indicates the lowest number in the series and orange indicates the highest number.

One way to consider these results is to address the transition point, or the point at which at least 50% of the experiments results in 70% or more of the phages having fitness-increasing mutations. If the increase in infectivity was set to 1 (*s*_*β*_=1); the transition point appears when the mutation opportunity was set to 5.0 × 10^−6^, (10 possible nucleotide sites) and the burst size increase for a mutation was set to 5-fold (Figure 7). The upper limit (all experiments having 70% mutations after four days) was reached when the mutation opportunity (U) was set to 1.0 × 10^−5^ (20 nucleotide sites) and burst size increased by 30-fold.

If the increase in infectivity in the event of a mutation was set to 5-fold; the transition point appears when the mutation opportunity (U) was set to 5.0 × 10^−7^ (1 nucleotide site) and burst size for a mutation was set to 10-fold (Figure 7). In this case, a single mutation would increase fitness by 50-fold. The upper limit was reached when the mutation opportunity (U) was set to 7.5 × 10^−6^ (15 nucleotide sites) and the burst size increase for a mutation was set to 5-fold, these settings designated an increase in fitness by 25-fold in the event of a single mutation.

The last set of simulations had the increase in infectivity in the event of a mutation set to 10-fold; the transition point in this instance appears when the mutation opportunity (U) was set to 5.0 × 10^−7^ (1 nucleotide site) and the burst size increase for the same mutation was set to 5-fold increase above the basal burst size of 10 (Figure 7). This effectively increased fitness by 50-fold if the mutation of that single available site occurred. The upper limit was reached when the mutation opportunity (U) was set to 5.0 × 10^−6^ (10 nucleotide sites) and the burst size increase for a mutation in one of these 10 sites was set to 5-fold increase in burst size; these mutations would increase overall fitness by 50-fold.

## 4. Discussion

Like many other phage biologists before us, we discovered *P. larvae* bacteriophages isolated from nature and found them to be intractable, meaning they had persistently low titers that could not be increased by standard laboratory procedures. Other *Paenibacillus* phage biologists have had similar issues. Yost and colleagues, working in the USA, described concentrating 100 mL volumes to 3 mL in order to obtain lysates for DNA extraction and EM (Yost et al., 2016). Similarly, a *P. polymyxa* bacteriophage isolated in Slovakia required the concentration of a 200 mL volume of lysate (Halgasova et al., 2010). More recently a similar study from Poland described five *P. larvae* bacteriophages, four of which were at a titer of 8×10^6^ or lower and required additional centrifugation steps before DNA sequencing (Jończyk-Matysiak et al., 2021). Similar protocols for tractable bacteriophages often require volumes of 1-10 mL and are easily brought to titers above 5×10^9^ PFU/mL.

We endeavoured to evolve one of our New Zealand *P. larvae* bacteriophages Lilo using traditional agar overlay methods, a 4.1 to 14.5 fold increase in titer was observed across the different replicates in the first 15 days; this increase was maintained for the rest of the duration of the experiment. We also saw increases in plaque size between Day 0 and Day 15 and then saw decreases in plaque size between Day 15 and Day 25. Increases in plaque size are complicated and can be due to a number of different factors. For example, shorter bacteriophage latent periods and faster virion diffusion can both lead to larger plaques sizes as well as a larger bacteriophage burst size, especially if initial burst size is small (Abedon, 2021). The increases in titer observed were not however sufficient to obtain enough DNA to progress to complete sequenced genomes.

Spatial structure could be a contributing factor to our difficulty in evolution of bacteriophages on agar overlay. There are many consequences of spatial structure including resource concentration, barriers and gradients, super-infection and altered gene expression (Bull et al., 2018). It is possible that gene expression changes in the host may determine which method works best for a particular bacteriophage (Chapman-McQuiston & Wu, 2008a, 2008b). In addition, the limits to diffusion that are in play when a phage is infecting in an agar overlay are significant.

We subsequently attempted the evolution of six New Zealand *P. larvae* bacteriophages using a modified Appelmans method, naturally evolving each bacteriophage independently on the bacterial strain on which they were isolated (Burrowes et al., 2019). This method resulted in a 100-fold increase in titer on average across the different bacteriophages. We also saw an increase in plaque size of between 63% and 170% in four of our bacteriophages. Interestingly, we saw the greatest increase in titer at approximately 4-5 days after commencing evolution. We therefore modified the protocol again and evolved a further six *P. larvae* bacteriophages using this new RAP method. From this protocol, we saw an average 1000-fold increase in titer and increases in plaque size ranging from 53% to 514%. The RAP method highlighted the possibility of evolving bacteriophages in even shorter timeframes and still obtaining desirable results.

The RAP described appears to facilitate mutations and the increase of these mutants in the population, as indicated by changes in titer and plaque dimensions. We sequenced and analysed the genomes of these experimentally evolved bacteriophages but couldn’t analyse their ancestors to determine what mutations led to the observed improvements. We did look at possible changes in CRISPR spaces that might have driven large increases in phage fitness by avoiding bacterial defence (Levin et al., 2013) but we could not find evidence that mutational escape from CRISPR drove these mutants.

Our experience of the RAP protocol and the increased titers of the phages did make us question whether it was possible that mutations could arise so quickly. We have therefore constructed a computer simulation of the population parameters of the RAP in order to determine whether advantageous mutations (and not some other phenomenon) could drive new phages to become dominant in the population in as little as four days.

We varied the target size for a random mutational event by changing the mutation opportunity (U) from 0.0 × 10^0^ to 1.0 × 10^−5^. We allowed single mutations to increase the infectivity from 1 to 10-fold and allowed the burst size to increase from 1 to 60-fold. Ultimately, we observed that with a mutation that would increase the fitness by only 5-fold as many as 50% of experiments would have at least 70% mutant phages. This phenomenon is likely similar to that of the mutational jackpot events, where beneficial mutations may occur early on in growth experiments resulting in these mutants dominating the population (Fusco et al., 2016; Luria & Delbrück, 1943). These results lead us to believe that this method may be highly relevant for increasing the utility of a wide range of difficult phages.

Serial passage experiments have previously been used to evolve bacteriophages. An elegant experiment to evolve four bacteriophages infecting *Pseudomonas aeruginosa* PAO1 in liquid media has been reported. At the end of the experiment, two of the bacteriophage isolates increased their infection capacity from 80-85% to 100% (Betts et al., 2013). In another experiment, bacteriophage FCV-1 was coevolved with *Flavobacterium columnare* in lake water, in the first four days the bacteriophage titer increased from 10^4^ PFU/mL to 10^7^ PFU/mL (approximately three log increase) (Laanto et al., 2020). The method that we have put forward here is a generalisable and high-throughput method that can be attempted rapidly with bacteriophages that are recalcitrant to other methods for amplification.

## 5. Conclusions

Communication with phage-hunting colleagues suggests that many bacteriophages have been discovered only to be abandoned due to issues with low titer. Herein, we have presented a novel method to experimentally evolve difficult phages to increase their titer and in some cases their host range in as little as four days. We have based this rapid method on the previously published Appelmans protocol (Burrowes et al., 2019). We have used this RAP to evolve 26 New Zealand *P. larvae* bacteriophages to rapidly improve stocks that are recalcitrant to high titer lysate creation by normal means. These high-titer lysates enable the extraction of high-quality DNA for sequencing as well as other downstream applications such as large-scale bacteriophage production. RAP presents a fast and effective way to experimentally evolve previously intractable phages allowing researchers to study entities that might otherwise be lost to science.

## Author Contributions

Conceptualization, H.L.H. and D.K.; methodology, D.K.; software, P.T., N.T.; validation, P.T., formal analysis, D.K.; investigation, J.T., D.K.; data curation, P.T.; writing— original draft preparation, D.K.; writing—review and editing, D.K., J.T., H.L.H., N.T., P.T.; visualization, D.K.; supervision, H.L.H.; project administration, H.L.H.; funding acquisition, H.L.H. All authors have read and agreed to the published version of the manuscript.

## Acknowledgements

Phage hunters everywhere.

## References

Abedon, S. T. (2016). Phage therapy dosing: The problem(s) with multiplicity of infection (MOI). Bacteriophage, 6(3), e1220348.

Abedon, S. T. (2021). Detection of Bacteriophages: Phage Plaques. In D. R. Harper, S. T. Abedon, B. H. Burrowes, & M. L. McConville (Eds.), Bacteriophages: Biology, Technology, Therapy (pp. 507–538). Springer International Publishing.

Abràmoff, & Magalhães. (n.d.). Image processing with ImageJ. Biophotonics International. https://dspace.library.uu.nl/handle/1874/204900

Betts, A., Vasse, M., Kaltz, O., & Hochberg, M. E. (2013). Back to the future: evolving bacteriophages to increase their effectiveness against the pathogen Pseudomonas aeruginosa PAO1. Evolutionary Applications, 6(7), 1054–1063.

Bonilla, N., Rojas, M. I., Netto Flores Cruz, G., Hung, S.-H., Rohwer, F., & Barr, J. J. (2016). Phage on tap-a quick and efficient protocol for the preparation of bacteriophage laboratory stocks. PeerJ, 4, e2261.

Brady, T. S., Merrill, B. D., Hilton, J. A., Payne, A. M., Stephenson, M. B., & Hope, S. (2017). Bacteriophages as an alternative to conventional antibiotic use for the prevention or treatment of Paenibacillus larvae in honeybee hives. Journal of Invertebrate Pathology, 150, 94–100.

Bull, J. J., Christensen, K. A., Scott, C., Jack, B. R., Crandall, C. J., & Krone, S. M. (2018). Phage-Bacterial Dynamics with Spatial Structure: Self Organization around Phage Sinks Can Promote Increased Cell Densities. Antibiotics (Basel, Switzerland), 7(1). https://doi.org/10.3390/antibiotics7010008

Burrowes, B. H., Molineux, I. J., & Fralick, J. A. (2019). Directed in Vitro Evolution of Therapeutic Bacteriophages: The Appelmans Protocol. Viruses, 11(3). https://doi.org/10.3390/v11030241

Chapman-McQuiston, E., & Wu, X. L. (2008a). Stochastic receptor expression allows sensitive bacteria to evade phage attack. Part I: experiments. Biophysical Journal, 94(11), 4525–4536.

Chapman-McQuiston, E., & Wu, X. L. (2008b). Stochastic receptor expression allows sensitive bacteria to evade phage attack. Part II: theoretical analyses. Biophysical Journal, 94(11), 4537–4548.

Cresawn, S. G., Bogel, M., Day, N., Jacobs-Sera, D., Hendrix, R. W., & Hatfull, G. F. (2011). Phamerator: a bioinformatic tool for comparative bacteriophage genomics. BMC Bioinformatics, 12, 395.

Delbrück, M. (1940). THE GROWTH OF BACTERIOPHAGE AND LYSIS OF THE HOST. The Journal of General Physiology, 23(5), 643–660.

Drake, J. W. (1991). A constant rate of spontaneous mutation in DNA-based microbes. Proceedings of the National Academy of Sciences of the United States of America, 88(16), 7160–7164.

Elke Genersch. (2010). American Foulbrood in honeybees and its causative agent, Paenibacillus larvae. Journal of Invertebrate Pathology, 103(S), S10–S19.

Ellis, E. L., & Delbrück, M. (1939). THE GROWTH OF BACTERIOPHAGE. The Journal of General Physiology, 22(3), 365–384.

Fusco, D., Gralka, M., Kayser, J., Anderson, A., & Hallatschek, O. (2016). Excess of mutational jackpot events in expanding populations revealed by spatial Luria–Delbrück experiments. Nature Communications, 7(1), 1–9.

Gallet, R., Kannoly, S., & Wang, I.-N. (2011). Effects of bacteriophage traits on plaque formation. BMC Microbiology, 11, 181.

Grissa, I., Vergnaud, G., & Pourcel, C. (2007). CRISPRFinder: a web tool to identify clustered regularly interspaced short palindromic repeats. Nucleic Acids Research, 35(Web Server issue), W52–7.

Halgasova, N., Ugorcakova, J., Gerova, M., Timko, J., & Bukovska, G. (2010). Isolation and characterization of bacteriophage PhiBP from Paenibacillus polymyxa CCM 7400. FEMS Microbiology Letters, 305(2), 128–135.

Hanauer, D. I., Graham, M. J., Sea-Phages, Betancur L., Bobrownicki, A., Cresawn, S. G., Garlena, R. A., Jacobs-Sera, D., Kaufmann, N., Pope, W. H., Russell, D. A., Jacobs, W. R., Jr, Sivanathan, V., Asai, D. J., & Hatfull, G. F. (2017). An inclusive Research Education Community (iREC): Impact of the SEA-PHAGES program on research outcomes and student learning. Proceedings of the National Academy of Sciences of the United States of America, 114(51), 13531–13536.

Hendrix, R. W., Smith, M. C., Burns, R. N., Ford, M. E., & Hatfull, G. F. (1999). Evolutionary relationships among diverse bacteriophages and prophages: all the world’s a phage. Proceedings of the National Academy of Sciences of the United States of America, 96(5), 2192–2197.

Jończyk-Matysiak, E., Owczarek, B., Popiela, E., Świtała-Jeleń, K., Migdał, P., Cieślik, M., Łodej, N., Kula, D., Neuberg, J., Hodyra-Stefaniak, K., Kaszowska, M., Orwat, F., Bagińska, N., Mucha, A., Belter, A., Skupińska, M., Bubak, B., Fortuna, W., Letkiewicz, S., … Górski, A. (2021). Isolation and Characterization of Phages Active against Paenibacillus larvae Causing American Foulbrood in Honeybees in Poland. Viruses, 13(7). https://doi.org/10.3390/v13071217

Laanto, E., Mäkelä, K., Hoikkala, V., Ravantti, J. J., & Sundberg, L.-R. (2020). Adapting a Phage to Combat Phage Resistance. Antibiotics (Basel, Switzerland), 9(6). https://doi.org/10.3390/antibiotics9060291

Lazeroff, M., Ryder, G., Harris, S. L., & Tsourkas, P. K. (2021). Phage Commander, an Application for Rapid Gene Identification in Bacteriophage Genomes Using Multiple Programs. PHAGE (New Rochelle, N.Y.), 2(4), 204–213.

LeMieux, J., & Hatfull, G. (2020). Set Phages to Kill: An Interview with Graham Hatfull, PhD. PHAGE (New Rochelle, N.Y.), 1(1), 4–9.

Levin, B. R., Moineau, S., Bushman, M., & Barrangou, R. (2013). The Population and Evolutionary Dynamics of Phage and Bacteria with CRISPR–Mediated Immunity. PLoS Genetics, 9(3), e1003312.

Luria, S. E., & Delbrück, M. (1943). Mutations of Bacteria from Virus Sensitivity to Virus Resistance. Genetics, 28(6), 491–511.

Mushegian, A. R. (2020). Are There 1031 Virus Particles on Earth, or More, or Fewer? Journal of Bacteriology, 202(9). https://doi.org/10.1128/JB.00052-20

Philipson, C. W., Voegtly, L. J., Lueder, M. R., Long, K. A., Rice, G. K., Frey, K. G., Biswas, B., Cer, R. Z., Hamilton, T., & Bishop-Lilly, K. A. (2018). Characterizing Phage Genomes for Therapeutic Applications. Viruses, 10(4). https://doi.org/10.3390/v10040188

Rohwer, F. (2003). Global phage diversity. Cell, 113(2), 141.

Russell, D. A., & Hatfull, G. F. (2017). PhagesDB: the actinobacteriophage database. Bioinformatics, 33(5), 784–786.

Salisbury, A., & Tsourkas, P. K. (2019). A Method for Improving the Accuracy and Efficiency of Bacteriophage Genome Annotation. International Journal of Molecular Sciences, 20(14). https://doi.org/10.3390/ijms20143391

Santos, M. A. (1991). An improved method for the small scale preparation of bacteriophage DNA based on phage precipitation by zinc chloride. Nucleic Acids Research, 19(19), 5442.

Skurnik, M., Pajunen, M., & Kiljunen, S. (2007). Biotechnological challenges of phage therapy. Biotechnology Letters, 29(7), 995–1003.

Stamereilers, C., Fajardo, C., Walker, J., Mendez, K., Castro-Nallar, E., Grose, J., Hope, S., & Tsourkas, P. (2018). Genomic Analysis of 48 Paenibacillus larvae Bacteriophages. Viruses, 10(7), 377–329.

Stamereilers, C., Wong, S., & Tsourkas, P. K. (2021). Characterization of CRISPR Spacer and Protospacer Sequences in Paenibacillus larvae and Its Bacteriophages. Viruses, 13(3). https://doi.org/10.3390/v13030459

Yost, D. G., Tsourkas, P., & Amy, P. S. (2016). Experimental bacteriophage treatment of honeybees (Apis mellifera) infected with Paenibacillus larvae, the causative agent of American Foulbrood Disease. Bacteriophage, 6(1), e1122698.

